# Phases of cortical actomyosin dynamics coupled to the neuroblast polarity cycle

**DOI:** 10.1101/2021.01.14.426740

**Authors:** Chet Huan Oon, Kenneth E. Prehoda

## Abstract

The Par complex dynamically polarizes to the apical cortex of asymmetrically dividing *Drosophila* neuroblasts where it directs fate determinant segregation. Previously we showed that apically directed cortical movements that polarize the Par complex require F-actin (Oon and Prehoda, 2019). Here we report the discovery of cortical actomyosin dynamics that begin in interphase when the Par complex is cytoplasmic but ultimately become tightly coupled to cortical Par dynamics. Interphase cortical actomyosin dynamics are unoriented and pulsatile but rapidly become sustained and apically-directed in early mitosis when the Par protein aPKC accumulates on the cortex. Apical actomyosin flows drive the coalescence of aPKC into an apical cap that is depolarized in anaphase when the flow reverses direction. Together with the previously characterized role of anaphase flows in specifying daughter cell size asymmetry, our results indicate that multiple phases of cortical actomyosin dynamics regulate asymmetric cell division.

## Introduction

The Par complex polarizes animal cells by excluding specific factors from the Par cortical domain (Lang and Munro, 2017; Venkei and Yamashita, 2018). In *Drosophila* neuroblasts, for example, the Par domain forms at the apical cortex during mitosis where it prevents the accumulation of neuronal fate determinants, effectively restricting them to the basal cortex. The resulting cortical domains are bisected by the cleavage furrow leading to fate determinant segregation into the basal daughter cell where they promote differentiation (Homem and Knoblich, 2012). It was recently discovered that apical Par polarization in the neuroblast is a multistep process in which the complex is initially targeted to the apical hemisphere early in mitosis where it forms a discontinuous meshwork (Kono et al., 2019; Oon and Prehoda, 2019). Cortical Par proteins then move along the cortex towards the apical pole, ultimately leading to formation of an apical cap that is maintained until shortly after anaphase onset (Oon and Prehoda, 2019). Here we examine how the cortical movements that initiate and potentially maintain neuroblast Par polarity are generated.

An intact actin cytoskeleton is known to be required for the movements that polarize Par proteins to the neuroblast apical cortex, but its role in the process has been unclear. Depolymerization of F-actin causes apical aPKC to spread to the basal cortex (Hannaford et al., 2018; Oon and Prehoda, 2019), prevents aPKC coalescence, and induces disassembly of the apical aPKC cap (Oon and Prehoda, 2019), suggesting that actin filaments are important for both apical polarity initiation and its maintenance. How the actin cytoskeleton participates in polarizing the Par complex in neuroblasts has been unclear, but actomyosin plays a central role in generating the anterior Par cortical domain in the *C. elegans* zygote. Contractions oriented towards the anterior pole transport the Par complex from an evenly distributed state (Illukkumbura et al., 2019; Lang and Munro, 2017). Bulk transport is mediated by advective flows generated by highly dynamic, transient actomyosin accumulations on the cell cortex (Goehring et al., 2011). While cortical movements of actomyosin drive formation of the Par domain in the worm zygote and F-actin is required for neuroblast apical Par polarity, no apically-directed cortical actomyosin dynamics have been observed during the neuroblast polarization process, despite extensive examination (Barros et al., 2003; Cabernard et al., 2010; Connell et al., 2011; Koe et al., 2018; Roth et al., 2015; Roubinet et al., 2017; Tsankova et al., 2017). Instead, both F-actin and myosin II have been reported to be cytoplasmic or uniformly cortical in interphase, and apically enriched at metaphase (Barros et al., 2003; Koe et al., 2018; Tsankova et al., 2017), before undergoing cortical flows towards the cleavage furrow that are important for cell size asymmetry (Cabernard et al., 2010; Connell et al., 2011; Roubinet et al., 2017).

The current model for neuroblast actomyosin dynamics is primarily based on the analysis of fixed cells or by imaging a small number of medial sections in live imaging experiments and we recently found that rapid imaging of the full neuroblast volume can reveal dynamic phases of movement that are not detected with other methods (LaFoya and Prehoda, 2021; Oon and Prehoda, 2019). Here we use rapid full volume imaging to investigate whether cortical actomyosin dynamics are present in neuroblasts when the Par complex undergoes its polarity cycle.

## Results and Discussion

### Cortical actin dynamics during the neuroblast polarity cycle

We imaged larval brain neuroblasts expressing an mRuby fusion of the actin sensor LifeAct (mRuby-LA) using spinning disk confocal microscopy. The localization of this sensor in neuroblasts has been reported (Abeysundara et al., 2018; Roubinet et al., 2017), but only during late mitosis. To follow cortical actin dynamics across full asymmetric division cycles, we collected optical sections through the entire neuroblast volume (~40 0.5 μm sections) at 10 second intervals beginning in interphase and through at least one mitosis (Figure 1-figure supplement 1). Maximum intensity projections constructed from these data revealed localized actin enrichments on the cortex, some of which were highly dynamic (Figure 1 and Video 1). We observed three discrete phases of cortical actin dynamics that preceded the previously characterized basally-directed flows that occur in late anaphase (Roubinet et al., 2017).

**Figure 1.**
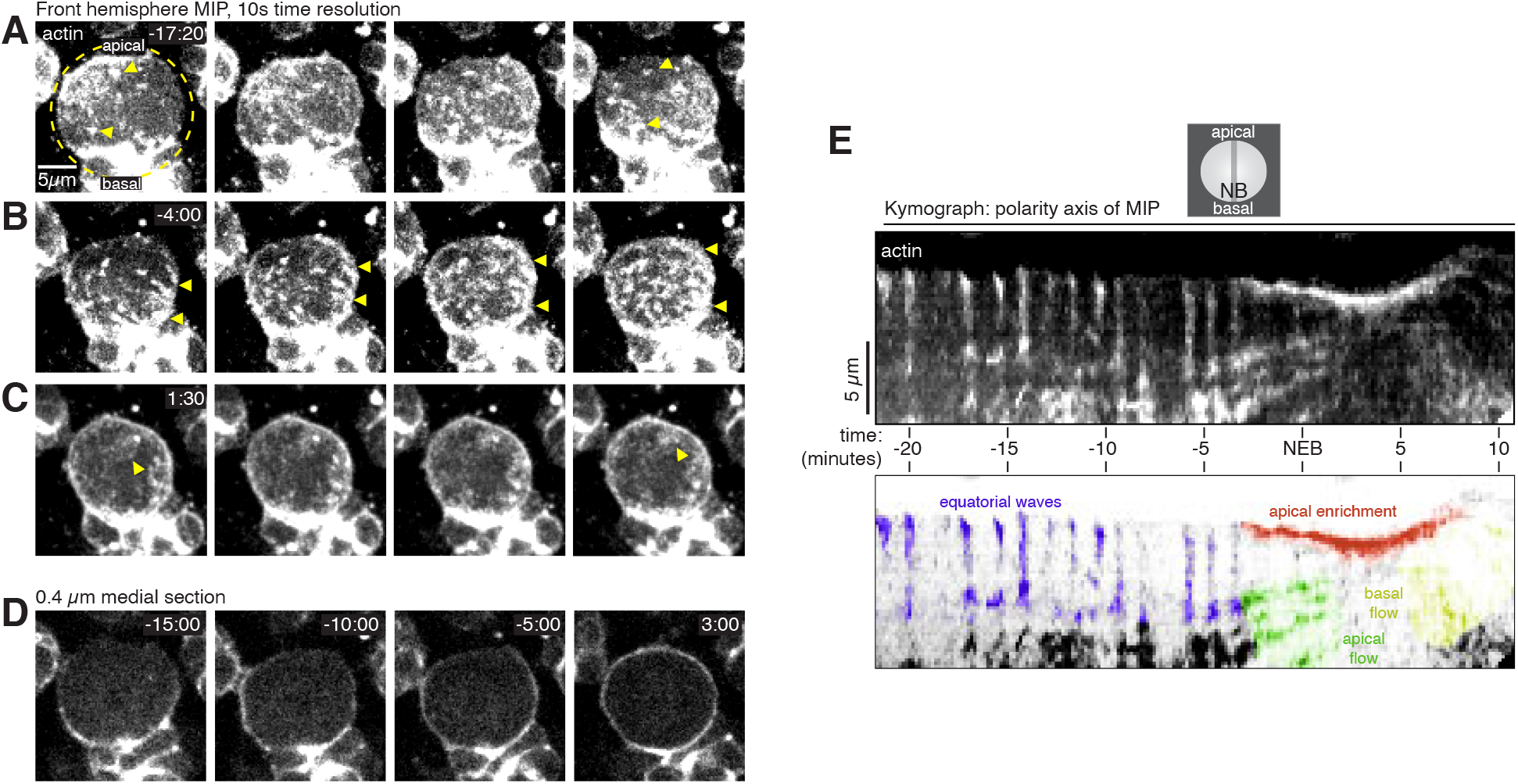
Cortical F-actin dynamics in asymmetrically dividing *Drosophila* larval brain neuro-blasts. (**A**) Selected frames from Video 1 showing cortical actin pulses during interphase. mRu-by-LifeAct expressed via insc-GAL4/UAS (“actin”) is shown via a maximum intensity projection (MIP) constructed from optical sections through the front hemisphere of the cell. The outline of the neuroblast is shown by a dashed yellow circle. In this example, the pulse moves from the upper left of the cell to the lower right. Arrowheads mark several cortical actin patches. Time (mm:ss) is relative to nuclear envelope breakdown. (**B**) Selected frames from Video 1 as in panel A showing cortical actin moving apically. Arrowheads delineate apical and basal extent of dynamic actin. (**C**) Selected frames from Video 1 as in panel A showing cortical actin enriched on the apical cortex. (**D**) Selected frames from Video 1 showing how actin becomes cortically enriched near NEB. Medial cross sections show the cortical actin signal is relatively discontinuous before NEB, with areas of verly low actin signal. (**E**) Kymograph constructed from frames of Video 1 using sections along the apical-basal axis as indicated. A legend for the features in the kymograph is included below.

The interphase neuroblast cortex was a mixture of patches of concentrated actin, highly dynamic pulsatile waves that traveled across the entire width of the cell, and areas with little to no detectable actin (Figure 1 and Video 1). Pulsatile movements consisted of irregular patches of actin forming on the cortex and rapidly moving across the surface before disappearing (Figure 1A,E). Concentrated actin patches were relatively static, but sometimes changed size over the course of several minutes. Static patches were mostly unaffected by the pulsatile waves that occasionally passed over them (Figure 1A). Pulses were sporadic in early interphase but became more regular near mitosis, with a new pulse appearing immediately following the completion of the prior one (Figure 1E and Video 1). The direction of the pulses during interphase was highly variable, but often along the cell’s equator (i.e. orthogonal to the polarity/division axis). In general, actin in the interphase cortex was highly discontinuous and included large areas with little to no detectable actin in addition to the patches and dynamic pulses described above (Figure 1D and Video 1). Interphase pulses were correlated with cellular scale morphological deformations in which these areas of low actin signal were displaced away from the cell center while the cortex containing the actin pulse was compressed towards the center of the cell (Figure 1D and Video 1).

Near nuclear envelope breakdown (NEB), the static cortical actin patches began disappearing from the cortex while dynamic cortical actin reoriented towards the apical pole. The unoriented and irregular pulses transitioned to more sustained, apically directed movements 3.4 ± 1.1 min before NEB (mean ± 1 SD; n = 13 neuroblasts from five larvae; Figure 1B-E and Video 1). In contrast to the sporadic and relatively unoriented interphase pulses observed earlier in the cell cycle, the apically-directed cortical actin dynamics that began near NEB were highly regular and were invariantly apically-directed (i.e. along the polarity/division axis; n = 13 neuroblasts from five larvae). This phase of cortical actin dynamics continued until anaphase–consistent with previous descriptions of actin accumulation at the apical cortex throughout metaphase (Barros et al., 2003; Tsankova et al., 2017). Additionally, while the interphase cortex had areas with very little actin, actin was more evenly-distributed following the transition, as was apparent in medial sections (e.g. comparing −15:00 and 3:00 in Figure 1D and Video 1).

Another transition in cortical actin dynamics occurred shortly after anaphase onset (7.3 ± 1.3 min after NEB; n = 13 neuroblasts from five larvae), when the apically-directed cortical actin movements rapidly reversed direction such that the F-actin that had accumulated in the apical hemisphere began to move basally towards the emerging cleavage furrow. The basally directed phase of movement that begins shortly after anaphase onset and includes both actin and myosin II was reported previously (Barros et al., 2003; Roubinet et al., 2017; Tsankova et al., 2017).

### Cortical actin flows mediate the aPKC polarity cycle

Previously we showed that Par polarity proteins undergo complex cortical dynamics during neuroblast asymmetric cell division and that polarity cycle movements require an intact actin cytoskeleton (Oon and Prehoda, 2019). Here we have found that the cortical actin cytoskeleton is also highly dynamic at points in the cell cycle when Par proteins undergo coordinated cortical movement (Figure 1 and Video 1). Furthermore, the transitions in cortical actin dynamic phases appeared to occur when similar transitions take place in the polarity cycle. We examined the extent to which cortical actin and aPKC dynamics are correlated by simultaneously imaging GFP-aPKC expressed from its endogenous promoter with mRuby-Lifeact (Figure 2 and Video 2). Apical targeting of aPKC began approximately ten minutes before NEB, when small, discontinuous aPKC foci began to appear, as previously reported (Oon and Prehoda, 2019). The interphase pulses of actin had no noticeable effect on these aPKC enrichments, suggesting that at this stage of the cell cycle, cortical actin dynamics are not coupled to aPKC movement (Figure 1C).

**Figure 2.**
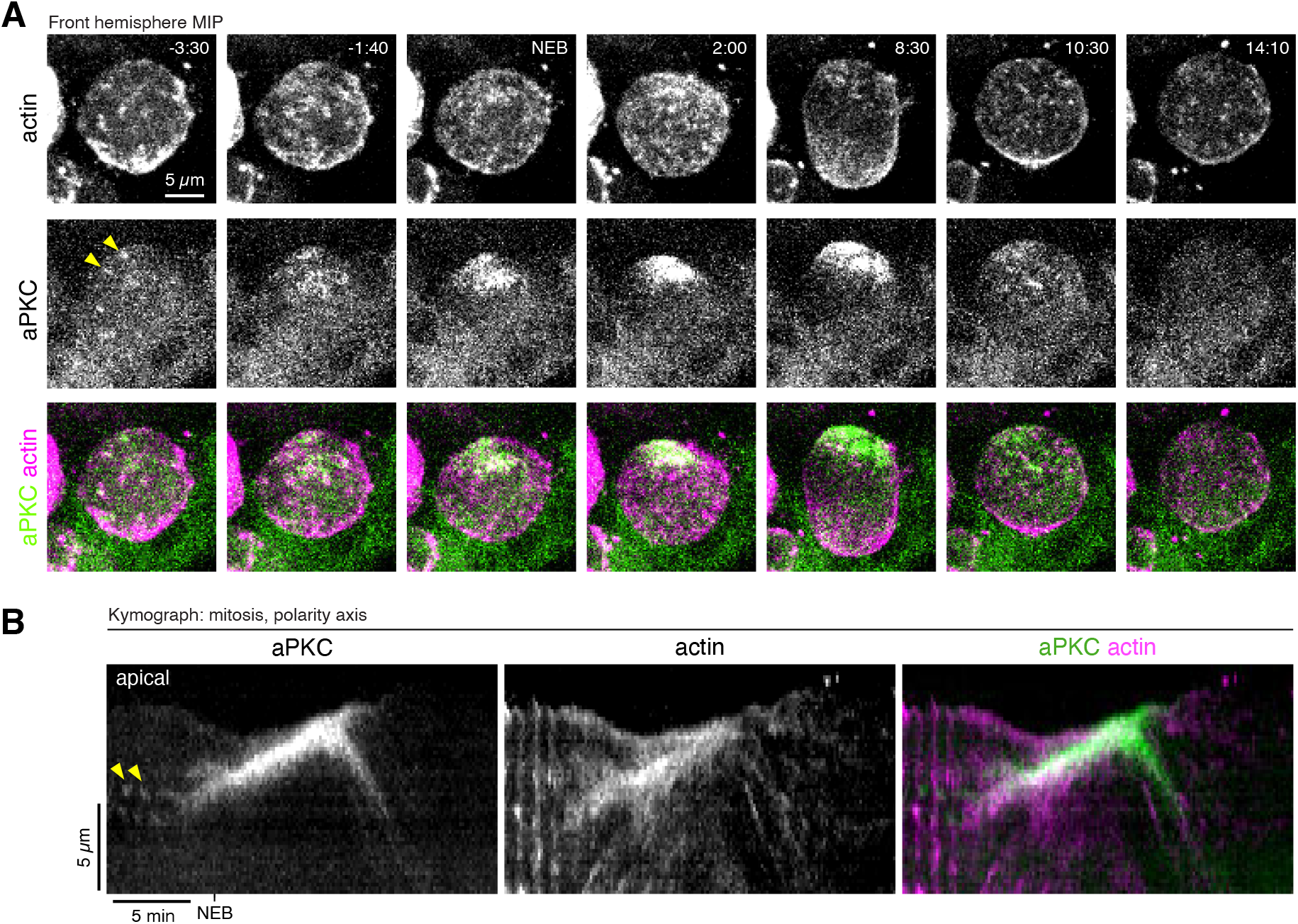
Coordinated actin and aPKC dynamics during the neuroblast polarity cycle. (**A**) Selected frames from Video 2 showing the correlated dynamics of aPKC and actin during polarization and depolarization. aPKC-GFP expressed from its endogenous promoter (“aPKC”) and mRu-by-LifeAct expressed via insc-GAL4/UAS (“actin”) are shown via a maximum intensity projection (MIP) constructed from optical sections through the front hemisphere of the cell. Arrowheads highlight several cortical aPKC foci. (**B**) Kymograph made from a segment along the apical-basal axis of the neuroblast in Video 2 showing the correlated dynamics of aPKC and actin. Arrowheads highlights cortical aPKC that is unaffected by equatorial actin pulses.

While cortical actin and aPKC did not appear to be coupled during interphase, the two protein’s movements were highly correlated beginning in early mitosis (Figure 2 and Video 2). When cortical actin began flowing apically the sparsely distributed aPKC patches that had accumulated on the cortex also began moving towards the apical pole. The transition to apically-directed movement was nearly simultaneous for both proteins, although actin’s apical movement began slightly before aPKC’s (actin: 1.9 ± 1.0 min prior to NEB; aPKC: 1.3 ± 1.1 min; n = 13 neuroblasts from 5 larvae). Furthermore, while aPKC and actin both moved towards the apical cortex, actin dynamics occurred over the entire cortex whereas aPKC movements were limited to the apical hemisphere consistent with its specific targeting to this area (Figure 2 and Video 2). The continuous apical movements resulted in the concentration of both aPKC and actin at the apical pole. Interestingly, however, once aPKC was collected near the pole into an apical cap, it became relatively static while cortical actin continued flowing apically. This phase of dynamic, apically-directed actin with an apparently static aPKC apical cap continued for several minutes (e.g. until approximately 8:30 in Video 2). At this point actin and aPKC movements reversed, moving simultaneously towards the basal pole and the emerging cleavage furrow (Figure 2 and Video 2). Thus, cortical actin and aPKC dynamics are highly correlated once the interphase actin pulses transition to sustained, apically-directed movements.

Simultaneous imaging cortical actin and aPKC allowed us to precisely determine when disruption of the actin cytoskeleton influences aPKC dynamics (Figure 3A,A’ and Figure 3-video 1). We introduced the actin depolymerizing drug Latrunculin A (LatA) and examined how the movement of cortical aPKC was influenced as the cortical actin signal dissipated. We previously found that the actin cytoskeleton is required for the apically-directed polarizing movements of aPKC (Oon and Prehoda, 2019). Here we find that when cortical actin is depolymerized during interphase (Figure 3A,A’ and Figure 3-video 1), aPKC is recruited to the apical cortex, but fails to undergo apically-directed movements and spreads prematurely to the basal pole (n = 36/36 neuroblasts from 15 larvae). Localized enrichments of aPKC have been reported at plasma membrane domains (LaFoya and Prehoda, 2021), and these domains normally move towards the apical pole shortly before metaphase. Consistent with previous observations (LaFoya and Prehoda, 2021), we find that these aPKC enrichments stop movement once cortical actin is depolymerized. Thus, cortical actin and aPKC dynamics are highly correlated during mitosis, and aPKC coalescence into an apical cap ceases immediately following the loss of cortical actin.

**Figure 3.**
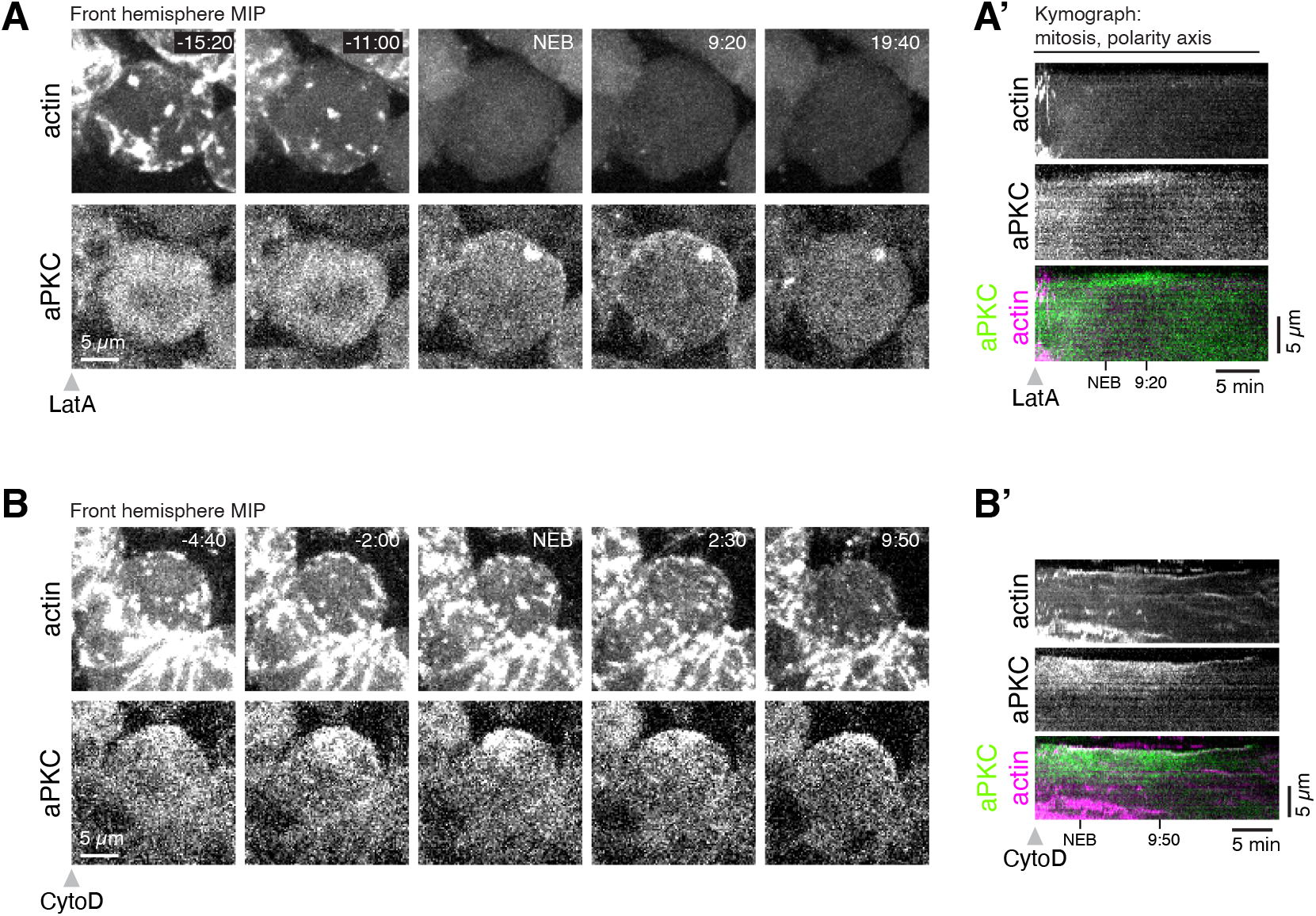
Cortical F-actin dynamics are required for aPKC coalescence. (**A**) Disruption of cortical F-actin with Latrunculin A causes immediate cessation of aPKC cortical movement. Selected frames from Figure 3-videos 1 showing how apically-directed aPKC movements cease upon complete loss of cortical actin. aPKC-GFP expressed from its endogenous promoter (“aPKC”) and mRuby-LifeAct expressed via insc-GAL4/UAS (“actin”) are shown via a maximum intensity projection (MIP) constructed from optical sections through the front hemisphere of the cell. (**A’**) Kymograph made from Figure 3-video 1 using a section of each frame along the apical-basal axis. (**B**) Selected frames from Figure 3-video 2 as in (A) showing how apically-directed aPKC movements cease upon loss of cortical actin dynamics using a low dosage (50 μM) of cytochalasin D. (**B’**) Kymograph made from Figure 3-video 2 using a section of each frame along the apical-basal axis.

We also examined the relationship between F-actin dynamics and aPKC using low doses of Cytochalasin D (CytoD) that inhibit actin dynamics without depolymerizing the filaments, thereby maintaining cortical structure (An et al., 2017; Mason et al., 2013). In these neuroblasts cortical F-actin signal persisted but did not undergo the apically-directed movements that occur in untreated neuroblasts (Figure 3B,B’ and Figure 3-video 2). The loss of cortical actin dynamics caused by CytoD was accompanied by relatively static cortical aPKC (we observed cessation of cortical actin dynamics in 29/36 neuroblasts from 13 treated larval brains and static aPKC in 29/29 neuroblasts). While aPKC coalescence was disrupted by the inhibition of cortical actin dynamics, aPKC was maintained in the apical hemisphere (n = 35/36 neuroblasts from 13 larvae; Figure 3B,B’ and Figure 3-Video 2), unlike in LatA treated neuroblasts. We conclude that cortical actin dynamics are required for aPKC coalescence into an apical cap.

### Myosin II dynamics are correlated with cortical actin

The morphological changes in interphase cells (Figure 1D and Video 1) and cortical aPKC movements that were correlated with cortical actin dynamics in early mitosis (Figure 2 and Video 2), are consistent with a force generating process. While actin can generate force directly through polymerization, contractile forces are generated by the combined activity of F-actin and myosin II (i.e. actomyosin), and cortical pulsatile contractions of actomyosin have been observed in many other systems (Vicker, 2000; Munro et al., 2004; Michaux et al., 2018). Although there have been several studies of myosin II dynamics in neuroblasts, no cortical dynamics have been reported and, like actin, its localization has been described as uniformly cortical or cytoplasmic in interphase and before metaphase in mitosis (Barros et al., 2003; Tsankova et al., 2017; Koe et al., 2018). We used rapid imaging of the full cell volume, simultaneously following a GFP fusion of the myosin II regulatory light chain Spaghetti squash (GFP-Sqh) with mRuby-Lifeact, to determine if myosin II undergoes cortical dynamics like actin. We found that myosin II is found with actin at every phase of cortical actin dynamics described above (Figure 4 and Video 3), including the apically-directed continuous movements that polarize aPKC. Interestingly, however, the localization between the two was not absolute and there were often large cortical regions where the two did not colocalize in addition to the region where they overlapped (Figure 4 and Video 3). A similar pattern of overlapping cortical actin and myosin II localization has been reported in the polarizing worm zygote (Reymann et al., 2016; Michaux et al., 2018). We conclude that there are multiple phases of cortical actomyosin dynamics during neuroblast asymmetric cell division.

**Figure 4.**
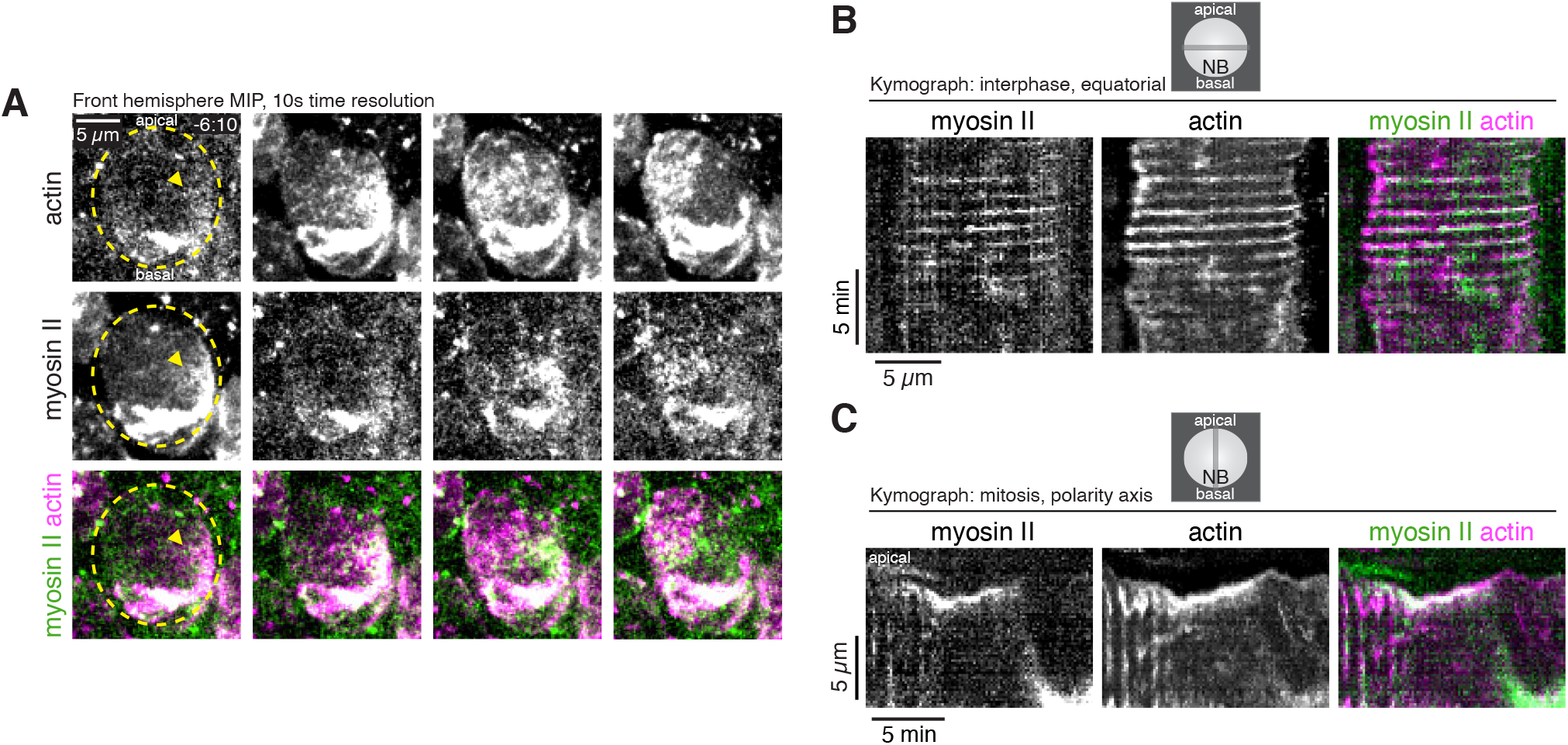
Dynamics of cortical actomyosin in asymmetrically dividing *Drosophila* larval brain neuroblasts. (**A**) Selected frames from Video 3 showing cortical actomyosin dynamics. GFP-Sqh expressed from its endogenous promoter (“Myosin II”) and mRuby-LifeAct expressed via wor-niu-GAL4/UAS (“actin”) are shown via a maximum intensity projection (MIP) constructed from optical sections through the front hemisphere of the cell. The outline of the neuroblast is shown by a dashed yellow line and arrowheads indicate the starting position of the cortical patches. Time is relative to nuclear envelope breakdown. (**B**) Kymograph constructed from frames of Video 3 during interphase using sections through the equatorial region of the cell as indicated. (**C**) Kymograph constructed from frames of Video 3 during mitosis using sections along the polarity axis of the cell as indicated.

### Phases of cortical actomyosin dynamics coupled to neuroblast polarization, maintenance, and depolarization

Our results reveal previously unrecognized phases of cortical actomyosin dynamics during cycles of neuroblast asymmetric divisions, several of which coincide with the neuroblast’s cortical polarity cycle (Figure 5). During interphase, transient cortical patches of actomyosin undergo highly dynamic movements in which they rapidly traverse the cell cortex, predominantly along the cell’s equator, before dissipating and beginning a new cycle (Figure 1A). Shortly after mitotic entry the movements become more continuous and aligned with the polarity axis (orthogonal to the equatorial interphase pulses). Importantly, the transition between these phases occurs shortly before the establishment of apical Par polarity, when discrete cortical patches of aPKC undergo coordinated movements towards the apical pole to form an apical cap. Apically-directed movements continue beyond metaphase when apical aPKC cap assembly is completed (Figure 2), suggesting that actomyosin dynamics may also be involved in cap maintenance. A role for actomyosin in aPKC cap assembly and maintenance is supported by the lack of coalescence when the actin cytoskeleton is completely depolymerized (Figure 3A,A’) or when actin dynamics are inhibited but the cytoskeleton is left intact (Figure 3B,B’). The cycle of cortical actomyosin dynamics is completed when the movement abruptly changes direction at anaphase leading to the cleavage furrow-directed flows that have been previously characterized (Barros et al., 2003; Roubinet et al., 2017).

**Figure 5.**
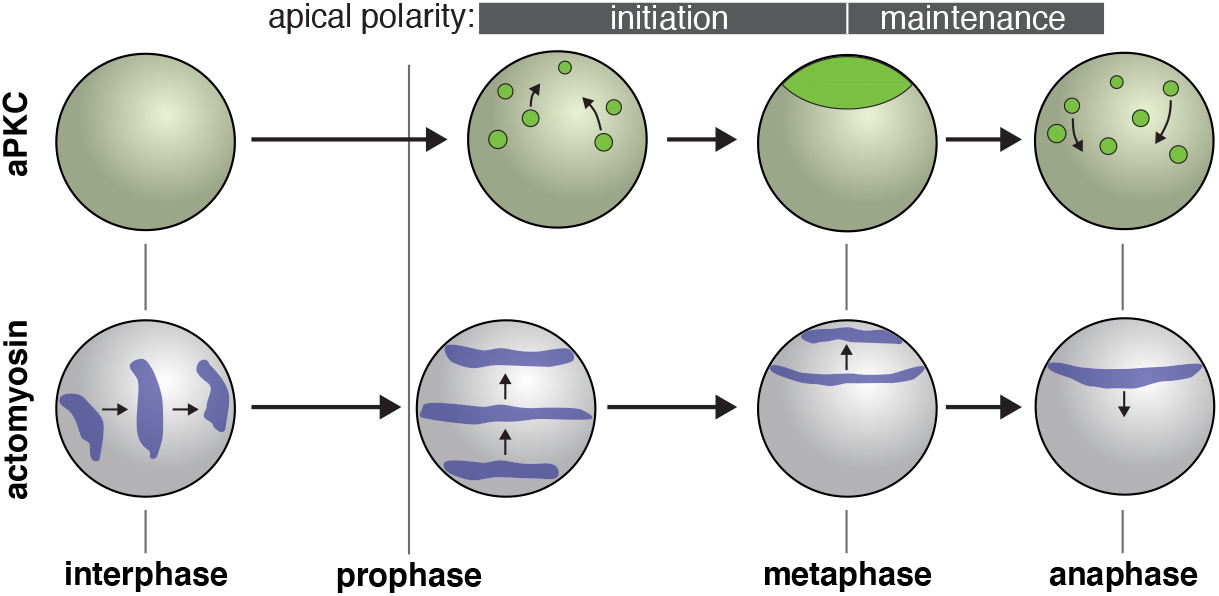
Model for role of actomyosin in neuroblast Par polarity. During interphase when aPKC is cytoplasmic, myosin II pulsatile contractions are predominantly equatorial. During apical polarity initiation in prophase and shortly before when discrete aPKC cortical patches begin to undergo coordinated movements towards the apical pole, myosin II pulsatile contractions reorient towards the apical cortex. Contractions are initially over a large surface area but become concentrated to the apical cortex as aPKC apical cap assembly is completed and the maintenance phase begins. At anaphase apical myosin II is cleared as it flows towards the cleavage furrow while the aPKC cap is disassembled.

While cortical actomyosin dynamics had not been reported during neuroblast polarization, myosin II pulses have been observed in delaminating neuroblasts from the *Drosophila* embryonic neuroectoderm (An et al., 2017; Simões et al., 2017). The actomyosin dynamics reported here may be related to those that occur during delamination and provide a framework for understanding how actomyosin participates in neuroblast apical polarity. First, apically directed pulsatile movements of actomyosin are consistent with the requirement for F-actin in the cortical flows that lead to coalescence of discrete aPKC patches (Figure 3) (Oon and Prehoda, 2019). Furthermore, the continuation of actomyosin pulsatile movements after cap assembly is completed (Figure 3), suggest a possible role in polarity maintenance. How might cortical actomyosin dynamics induce aPKC coalescence and maintenance? In the worm zygote, pulsatile contractions generate bulk cortical flows (i.e. advection) that lead to non-specific transport of cortically localized components (Goehring et al., 2011; Illukkumbura et al., 2019). Whether the cortical motions of polarity proteins that occur during the neuroblast polarity cycle are also driven by advection will require further study.

## Materials and Methods

### Key Resources Table

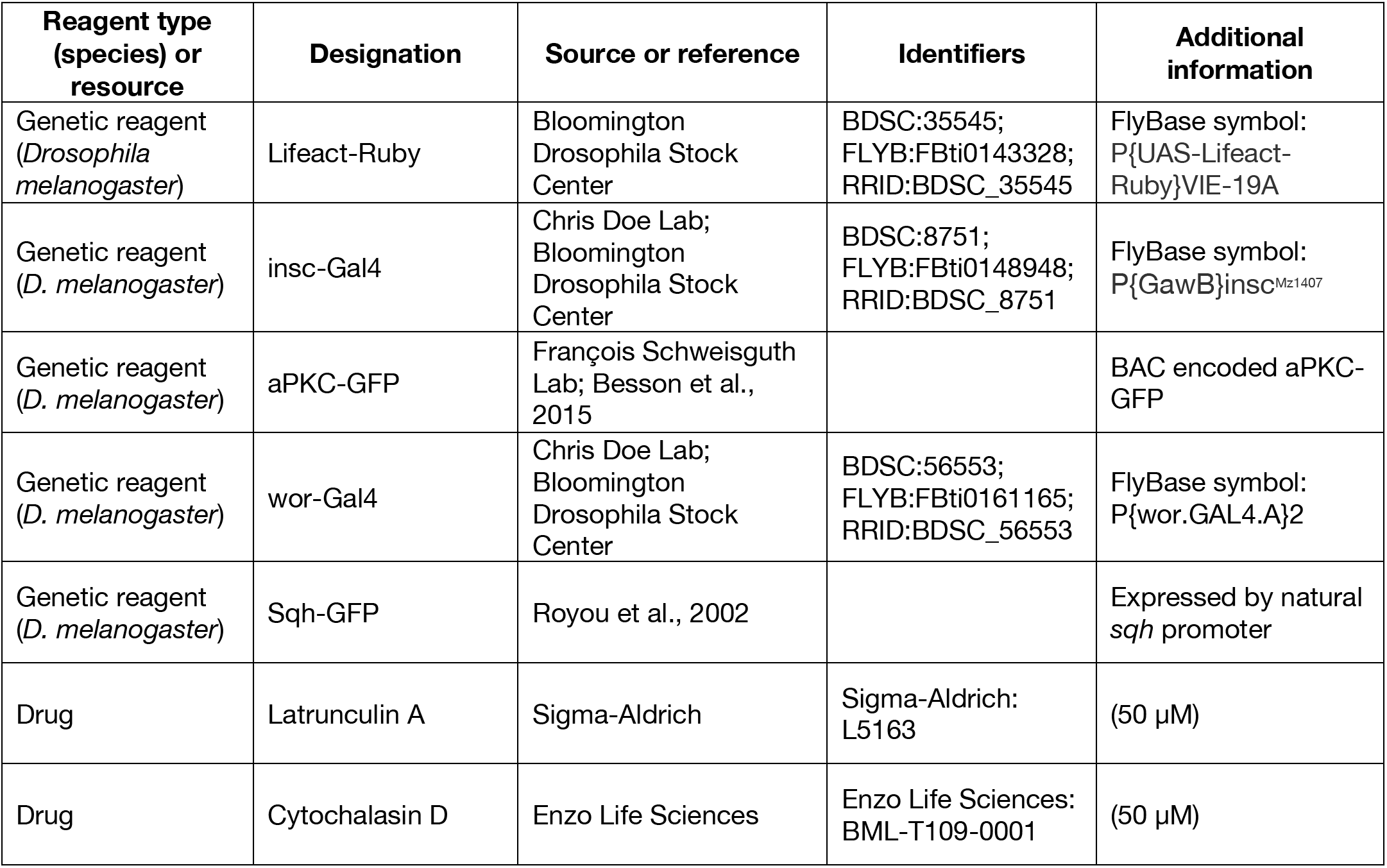

| Reagent type (species) or resource | Designation | Source or reference | Identifiers | Additional information |
| --- | --- | --- | --- | --- |
| Genetic reagent (*Drosophila melanogaster*) | Lifeact-Ruby | Bloomington Drosophila Stock Center | BDSC:35545; FLYB:FBti0143328; RRID:BDSC_35545 | FlyBase symbol: P{UAS-Lifeact-Ruby}VIE-19A |
| Genetic reagent (*D. melanogaster*) | insc-Gal4 | Chris Doe Lab; Bloomington Drosophila Stock Center | BDSC:8751; FLYB:FBti0148948; RRID:BDSC_8751 | FlyBase symbol: P{GawB}insc<sup>Mz1407</sup> |
| Genetic reagent (*D. melanogaster*) | aPKC-GFP | François Schweisguth Lab; Besson et al., 2015 |  | BAC encoded aPKC-GFP |
| Genetic reagent (*D. melanogaster*) | wor-Gal4 | Chris Doe Lab; Bloomington Drosophila Stock Center | BDSC:56553; FLYB:FBti0161165; RRID:BDSC_56553 | FlyBase symbol: P{wor.GAL4.A}2 |
| Genetic reagent (*D. melanogaster*) | Sqh-GFP | Royou et al., 2002 |  | Expressed by natural *sqh* promoter |
| Drug | Latrunculin A | Sigma-Aldrich | Sigma-Aldrich: L5163 | (50 μM) |
| Drug | Cytochalasin D | Enzo Life Sciences | Enzo Life Sciences: BML-T109-0001 | (50 μM) |

#### Fly strains and genetics

UAS-Lifeact-Ruby (Bloomington stock 35545), BAC-encoded aPKC-GFP (Besson et al., 2015) and Sqh-GFP (Royou et al., 2002) transgenes were used to assess F-actin, aPKC and myosin II dynamics, respectively. Expression of Lifeact was specifically driven in nerve cells upon crossing UAS-Lifeact-Ruby to *insc-Gal4* (1407-Gal4, Bloomington stock 8751) or to *worniu-Gal4* (Bloomington stock 56553). The following genotypes were examined through dual channel live imaging: BAC-aPKC-GFP / Y; insc-Gal4, + / +, UAS-Lifeact-Ruby; and; worGal4, Sqh-GFP, + / +, UAS-Lifeact-Ruby;.

#### Live imaging

Third instar larvae were incubated in 30°C overnight (~12 hours) prior to imaging and were dissected to isolate the brain lobes and ventral nerve cord, which were placed in Schneider’s Insect media (SIM). Larval brain explants were placed in lysine-coated 35 mm cover slip dishes (WPI) containing modified minimal hemolymph-like solution (HL3.1). Explants were imaged on a Nikon Ti2 microscope equipped with a Yokogawa CSU-W1 spinning disk that was configured to two identical Photometrics Prime BSI Scientific CMOS cameras for simultaneous dual channel live imaging. Using the 1.2 NA Plan Apo VC water immersion objective, explants were magnified at 60x for visualization. Explants expressing Lifeact-Ruby, aPKC-GFP and Sqh-GFP were illuminated with 488 nm and 561 nm laser light throughout approximately 41 optical sections with step size of 0.5 μm and time interval of 10 seconds. To observe the effects of dissipating F-actin and inhibiting F-actin dynamics on aPKC dynamics, explants expressing Lifeact-Ruby and aPKC-GFP were treated with 50 μM LatA (2% DMSO) and 50 μM CytoD (0.5% DMSO) prior to imaging, respectively.

#### Image processing, analysis and visualization

Movies were analyzed in ImageJ (using the FIJI package) and in Imaris (Bitplane). Neuroblasts whose apical-basal polarity axis is positioned parallel to the imaging plane were cropped out to generate representative images and movies. Cortical edge and central maximum intensity projections (MIP) were derived from optical slices capturing the surface and center of the cell, respectively. Cortical MIPs were also used to perform kymograph analysis, where the change in localization profile of fluorescently-tagged fusion proteins within a 3 to 5 pixels wide region was examined across time. To track cortical movements over the length of the apical-basal axis, a vertical region parallel to the polarity axis was specified for the kymograph analysis. Similarly, a horizontal line orthogonal to the polarity axis that is superimposing on the presumptive equator was specified for examining equatorial motions. Optical sections capturing the whole of the cell were assembled for 3D rendering and visualization in FIJI or Imaris. These volumetric reconstructions were then used to determine the timing of cortical motions characterized in this paper.

### Video Legends

Video 1 Actin dynamics in a larval brain neuroblast.

The mRuby-Lifeact sensor expressed from the UAS promoter and *insc-GAL4* (expressed in neuroblasts and their progeny) is shown with a maximum intensity projection of the front hemisphere of the cell.

Video 2 Correlated dynamics of the Par protein aPKC and actin in a larval brain neuroblast. GFP-aPKC expressed from its endogenous promoter and the mRuby-Lifeact sensor expressed from the UAS promoter and *insc-GAL4* (drives expression in neuroblasts and progeny) are shown from simultaneously acquired optical sections with a maximum intensity projection of the front hemisphere of the cell.

Figure 3-video 1 Correlated dynamics of the Par protein aPKC and actin in a larval brain neuroblast treated with Latrunculin A before mitosis.

GFP-aPKC expressed from its endogenous promoter and the mRuby-Lifeact sensor expressed from the UAS promoter and *insc-GAL4* (drives expression in neuroblasts and progeny) are shown from simultaneously acquired optical sections with a maximum intensity projection of the front hemisphere of the cell. LatA was added to the media surrounding the larval brain explant at the indicated time.

Figure 3-video 2 Correlated dynamics of the Par protein aPKC and actin in a larval brain neuroblast treated with Cytochalasin D during prophase.

GFP-aPKC expressed from its endogenous promoter and the mRuby-Lifeact sensor expressed from the UAS promoter and *insc-GAL4* (drives expression in neuroblasts and progeny) are shown from simultaneously acquired optical sections with a maximum intensity projection of the front hemisphere of the cell. CytoD was added to the media surrounding the larval brain explant at 2 minutes prior to the beginning of the movie.

Video 3 Correlated dynamics of myosin II and actin in a larval brain neuroblast.

GFP-Sqh (the myosin II regulatory light chain, Spaghetti squash) expressed from its endogenous promoter and the mRuby-Lifeact sensor expressed from the UAS promoter and *worniu-GAL4* (expressed in neuroblasts and their progeny) are shown from simultaneously acquired optical sections with a maximum intensity projection of the front hemisphere of the cell and the medial optical section. The neuroblast is highlighted by a dashed circle.

---

**Figure 1 – figure supplement 1.**
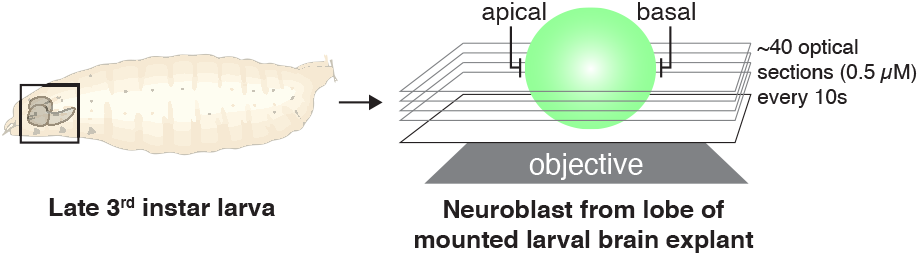
Imaging and analysis scheme for rapid, full volume imaging of *Drosophila* neuroblasts from larval brain explants. Larval brains from 3rd instar larvae were mounted and imaged along the neuroblast polarity axis (“apical” and “basal”). Optical sections across the full cell volume were acquired every 10 seconds and used to construct maximum intensity projections.

